# Beta-catenin mediates growth defects induced by centrosome loss in APC mutant colorectal cancer independently of p53

**DOI:** 10.1101/2023.02.13.528233

**Authors:** Mohamed Bourmoum, Nikolina Radulovich, Ming-Sound Tsao, Laurence Pelletier

## Abstract

Colorectal cancer is the third most common cancer and the second leading cause of cancer-related deaths worldwide. The centrosome is the main microtubule-organizing center in animal cells and centrosome amplification is a hallmark of cancer cells. To investigate the importance of centrosomes in colorectal cancer, we induced centrosome loss in normal and cancer human-derived colorectal organoids using centrinone B, a Polo-like kinase 4 (Plk4) inhibitor. We show that centrosome loss represses human normal colorectal organoid growth in a p53-dependent manner in accordance with previous studies in cell models [1]. However, cancer colorectal organoid lines exhibited different sensitivities to centrosome loss independently of p53. Centrinone-induced cancer organoid growth defect/death positively correlated with a loss of function mutation in the APC gene, suggesting a causal role of the hyperactive WNT pathway. Consistent with this notion, β-catenin inhibition using XAV-939 or ICG-001 partially prevented centrinone-induced death and rescued the growth of APC-mutant organoid lines. Our study reveals a novel role for canonical WNT signaling in regulating centrosome loss-induced growth defect/death in APC-mutant colorectal cancer independently of the classical p53 pathway.

## Introduction

Colorectal cancer (CRC) is the third most common and the second most deadly cancer type worldwide (almost 2 million cases and 1 million deaths in 2020) [2]. CRC is a highly heterogeneous disease. Large-scale transcriptomic analyses led to CRC classification into four consensus molecular subtypes (CMS): CMS1 (MSI Immune, 14%), hypermutated, microsatellite unstable, strong immune activation; CMS2 (Canonical, 37%), epithelial, chromosomally unstable, pronounced WNT and MYC signaling activation; CMS3 (Metabolic, 13%), epithelial, evident metabolic dysregulation; and CMS4 (Mesenchymal, 23%), marked TGF-β activation, stromal invasion, and angiogenesis [3]. Mutations affecting key genes regulating cell proliferation, differentiation, and death accumulate in neoplastic cells, giving them a survival advantage over surrounding normal intestinal epithelial cells [4]. These mutated genes lead to abnormal expansion of premalignant tissue into adenomas that have the potential to completely transform into invasive carcinomas due to additional genetic aberrations [4, 5]. Alterations in certain genes, such as APC and KRAS, have been shown to occur early whereas other genetic events are typically only observed in more advanced disease states, like the loss of function p53 mutations [4, 6].

The centrosome is the major microtubule-organizing centre (MTOC) in eukaryotic cells, consisting of two centrioles surrounded by an electron-dense matrix, the pericentriolar material (PCM). Centrosomes regulate many cellular processes including cell motility, adhesion, and polarity in interphase, and organizing the spindle assembly during mitosis [7]. In addition, in quiescent cells, centrioles can act as basal bodies that anchor cilia and flagella, which play important roles in physiology, development and disease [8]. Numerical and structural centrosome aberrations are frequent in many cancers and can be associated with genomic instability. Whether changes in centrosome numbers are a cause or consequence of oncogenic transformation remains a matter of debate [9] but in certain cancer types, evidence suggests that centrosomal defects might occur very early in tumorigenesis [10]. Centriole duplication is tightly controlled so that cells have precisely two centrosomes [11]. The serine-threonine protein kinase Polo-like kinase 4 (Plk4) plays a pivotal role in this process [12]. Wong et al [1] developed centrinone B, a selective Plk4 inhibitor and have found that centrinone-induced centrosome loss irreversibly arrested normal cells in a senescence-like G1 state by a p53-dependent mechanism whereas p53-mutant cancer cells could proliferate indefinitely after centrosome loss.

The development of human organoids has greatly benefited the biomedical research community as they filled the gap between the animal models, lacking human specificity, and the 2D cell models, lacking biological complexity Since their establishment, human colorectal organoids [13] have been widely used to address different biological questions and to model colorectal cancer.

Here, using centrinone B, we sought to determine the role of centrosomes in the growth and survival of normal and cancer human-derived colorectal organoids (HCOs).

## Results

### Centrosome loss represses human normal colon organoid growth in a p53-dependent manner

We first assessed the effect of centrosome depletion on the normal HCOs growth using centrinone B. Normal colon organoids were grown from single adult intestinal stem cells in the presence of DMSO or centrinone B. Centrinone B treatment strongly repressed the growth of the human normal organoids measured by the relative average organoid area (Fig1A, 1B) (~ 88% reduction of the average organoid area). Previous studies have shown that centrosome loss irreversibly arrested normal cells in a senescence-like G1 state by a p53-dependent mechanism [1]. To test this in our model, we generated a p53 knockout human colon organoid line by infecting the adult stem cells with an inducible lentiviral system expressing CRISPR/Cas9 and the p53 gRNA. p53-/- organoids were enriched by Nutlin-3a selection for 2 weeks as described previously [14]. INDELs at the p53 gene locus were confirmed using TIDE analysis [15] (Fig 1D) and the loss of p53 protein was confirmed by immunofluorescence (Fig 1E). We noticed that the p53-/- normal organoid growth rate was higher (~55% bigger average organoid area). More interestingly, we found that p53 depletion partially rescued the human organoid growth in the presence of centrinone B (~55% decrease in the average organoid area after centrinone B treatment in p53-/- organoids versus 88% decrease in wild-type organoids) (Fig1A, 1B) indicating that the centrinone-induced growth defect in human colon organoid is p53-dependent in accordance with previous reports in tissue culture cells [1]. In wild-type and p53-/- human normal colon organoids, centrinone-induced centrosome loss was confirmed by staining for the centrosomal protein CEP192 as shown in Fig 1C. It is important to mention that despite continuing to grow in the absence of centrosomes, p53-/- normal organoids exhibited recurrent abnormal nuclei phenotypes including big multi-lobed nuclei and multinucleated cells suggesting frequent mitotic defects (data not shown).

**Fig1.**
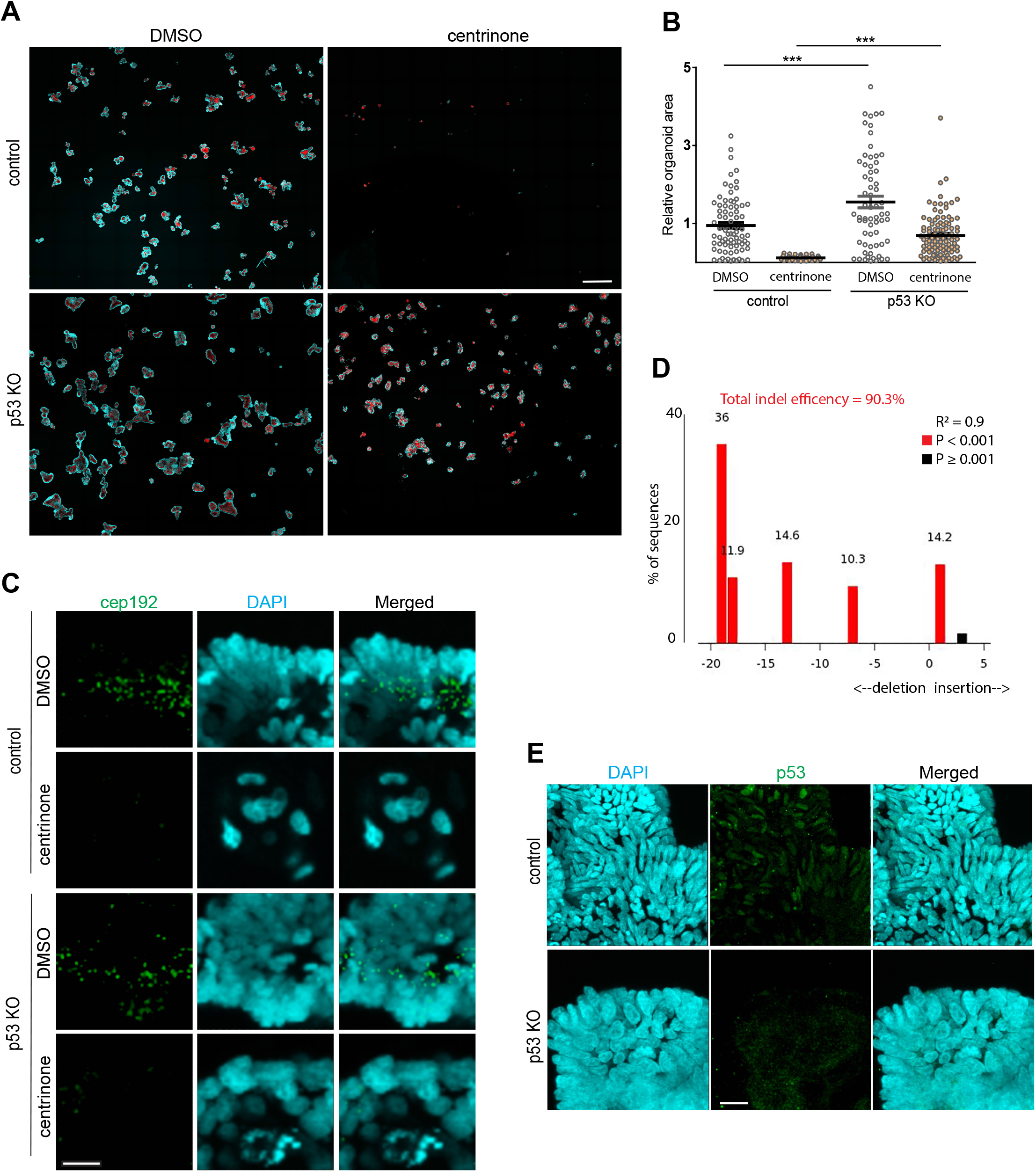
Centrosome loss represses normal HCOs growth in a p53-dependent manner. **A)** Control and p53 KO normal HCOs were grown from single adult stem cells in the presence of DMSO or 0.5 μM centrinone B for 14 days. Organoids were then fixed, stained with DAPI to label DNA (blue) and phalloidin to label actin (red) and imaged. Merged maximum intensity projections of images are shown. **B)** The areas of individual organoids were quantified in the merged maximum intensity projections (MIPs) of images (from A) (n=3) and presented in the graph as relative values, every dot represents an organoid (\*\*\**P*<0.001). **C)** High-resolution maximum intensity projection images of CEP192 (centrosome) and DNA (DAPI) immunofluorescence staining in selected normal HCOs from (A). **D)** Tide analysis of p53 gene indel efficiency in the normal HCO p53 KO line. The software provides the R2 value as a goodness-of-fit measure and calculates the statistical significance for each indel. Red represents significant indels (P<0.001) Note that the total indel efficiency is ~90.3%. **E)** Representative maximum intensity projections of p53 immunofluorescence staining in control and p53 KO normal HCOs. Scale bars are 500 μm (A), 10 μm (B) and 25 μm (C).

### Patient-derived cancer colorectal organoids exhibit different sensitivity to centrosome loss independently of p53

To determine whether centrosomes are essential for colorectal cancer growth, we next assessed the effect of centrosome depletion on patient-derived cancer HCOs growth similar to what we did with the normal HCOs. We performed centrosome loss experiments in three cancer colorectal organoid lines derived from three different patients (CSC-406, POP-092 and POP-112, see methods). In contrast to previous studies in cell models [1], we observed that the three cancer colorectal organoid lines exhibited different sensitivity to centrosome loss independently of p53. Whole exome sequencing data showed that all the colorectal cancer lines carried a missense p53 mutation (S240R in CSC-406, R248W in POP-092 and R248Q in POP-112) (Supplementary file1, Fig2D). These mutations are in the DNA-binding domain of p53 and are known to perturb its function as a tumor suppressor [16, 17]. Despite a nonfunctional p53, centrinone B treatment strongly repressed the growth of CSC-406 and POP-092 cancer colorectal organoids (~93% and 84% decrease in the average organoid area respectively) (Fig2A, 2B) whereas the POP-112 line was relatively resistant to centrosome loss (only ~22% decrease in the average organoid area) (Fig2A, 2B). To confirm that the centrinone-induced growth defect observed in CSC-406 and POP-092 lines was indeed p53-independent, we generated p53 null alleles in these lines using a lentiviral CRISPR-CAS9. p53 depletion efficiency was verified by western blot analysis (Fig 2C). Our results show that p53 Knockout did not prevent the centrinone-induced growth defect in both CSC-406 and POP-092 lines confirming that the latter is p53-independent (Fig2A, 2B). We then asked whether POP-112 resistance to centrinone-induced growth defect is due to a lower sensitivity to centrinone B. Immunofluorescence staining for pericentrin, a centrosomal marker, confirmed that the centrinone B dose used (0.5 μM) led to efficient centrosome depletion (Fig 2E). We further carried out dose-dependent growth experiments in the POP-112 line. As shown in Fig 2F, a 10 μM centrinone B concentration was needed to induce a growth defect (~95% decrease in the average organoid area) comparable to the extent of growth defect observed in CSC-406 and POP-092 with only 0.5 μM concentration. This dose (10 μM) is 20X higher than the working centrinone B dose (0.5 μM) needed to deplete centrosomes in the POP-112 line, suggesting a centrosome-independent mechanism inducing cell death.

**Fig2.**
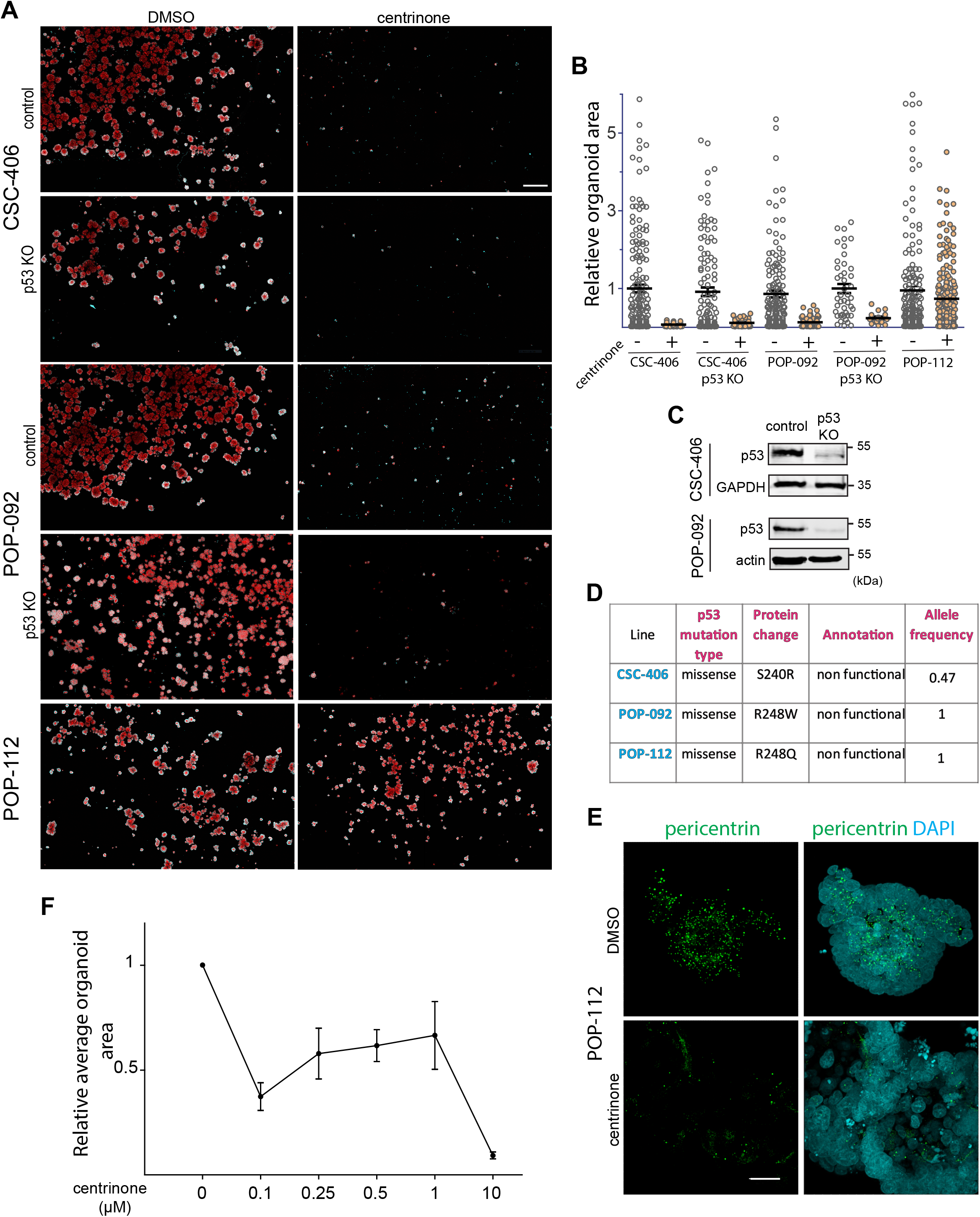
Patient-derived cancer HCOs exhibit different sensitivity to centrosome loss independently of p53. **A)** Cancer HCO lines (control or p53 KO) were grown from single adult stem cells in the presence of DMSO or 0.5 μM centrinone B for 8 days. Organoids were then fixed, stained with DAPI (blue) and phalloidin (red) and imaged. Merged maximum intensity projection images are shown. **B)** The areas of individual organoids were quantified in the merged maximum intensity projection images (from A) (n=3) and presented in the graph as relative values; every dot represents an organoid (\*\**P*<0.01\*\*\**P*<0.001). **C)** Western blot analysis of p53 and loading controls (GAPDH and actin) in lysates prepared from control and p53 KO Cancer HCO lines. **D)** p53 mutation status in the three cancer HCOs from the whole exome sequencing data. **E)** High-resolution maximum intensity projections of pericentrin (centrosome) and DNA (DAPI) immunofluorescence staining in POP-112 organoids treated with DMSO or 0.5 μM centrinone B for 8 days. **F**) POP-112 organoids were grown from single adult stem cells in the presence of DMSO (0) or the indicated centrinone B concentrations for 8 days. Organoid areas were quantified as in (B). Relative average organoid area from three independent experiments is presented in the graph (n=3). Scale bars are 500 μm (A) and 25 μm (E).

Altogether, our results show that centrosome loss inhibits human normal colon organoid growth in a p53-dependent manner. However, cancer colorectal organoids exhibit different sensitivity to centrosome loss independently of p53, suggesting an additional regulatory mechanism.

### Mechanisms underlying centrinone sensitivity in cancer HCOs

Our data show that the centrinone-induced growth defect observed in CSC-406 and POP-092 cancer lines is p53-independent (Fig 2). The three cancer organoid lines used were derived from different patients and are expected to carry different mutation signatures. To pinpoint the potential mechanisms involved in this growth defect, we first examined the mutation status of the main known colorectal cancer driver genes besides p53 (APC, K-Ras, SMAD4) [18]. Whole exome sequencing data (Supplementary file1, Fig 3A) revealed a positive correlation between sensitivity to centrinone B and the presence of non-functional APC alleles. Indeed, the two centrinone-sensitive lines CSC-406 and POP-092 carried a non-functional APC mutation (nonsense Q1045* / Fs insertion P1594Afs*38 mutations for CSC-406 and nonsense G1499* mutation for POP-092) whereas the centrinone-resistant line, POP-112 possessed wild-type APC. These results suggested that centrinone-induced organoid growth defect might be mediated by hyperactive WNT signaling. To test this hypothesis, we inhibited the WNT signaling pathway downstream of APC using XAV-939, a tankyrase inhibitor that increases the protein levels of axin, thereby promoting the degradation of β-catenin, the downstream effector of the canonical WNT pathway [19]. We treated CSC-406 organoids with increasing doses of XAV-939 (0,5,10 and 20 μM) in the presence or absence of centrinone B (0.5μM) (Fig 3B, C, D). We noted that concentrations above 20 μM of XAV-939 decreased organoid growth (data not shown). Organoid growth (average organoid area) and survival rate (relative number of surviving organoids) were then assessed. As shown in Fig 3C and 3D, centrinone B treatment resulted in a~ 85% decrease in surviving organoid number accompanied by a ~75% reduction in the organoid size. Remarkably, increasing doses of XAV-939 rescued the survival and the growth rate of centrinone B-treated organoids. A 20 μM XAV-939 treatment significantly prevented ~75% centrinone-induced organoid death and increased the growth rate of the surviving organoids by ~40% (Fig3 C, 3D). This XAV-939 concentration (20 μM) (we considered as the optimal one) efficiently inhibited the WNT pathway as it induced a ~67% decrease in β-catenin protein level as shown in western bot analysis (Fig 3E). To confirm our results, we targeted the WNT pathway using another inhibitor, Foscenvivint (ICG-001) that antagonizes β-catenin/TCF-mediated transcription by specifically binding to CREB-binding protein (CBP) [20]. We tested different ICG-001concentrations and found that 3 μM is the maximal dose relatively tolerated by CSC-406 cancer organoids as higher concentrations notably affected organoid growth and survival (data not shown). Our data (Fig3 F, G) show that, similarly to XAV939, ICG-001 partially prevented the centrinone-induced organoid death (~38 % rescue of organoid survival rate) but the effect on growth rate was non-significant (Fig 3H) probably because the 3 μM ICG-001 treatment also reduced the DMSO-treated CSC-406 organoid growth (~30% reduction in average organoid area) (Fig 3H). Unlike XAV-939 and ICG-001 which act downstream of the APC protein to inhibit the WNT pathway, using DKK1, a negative regulator of the WNT pathway that acts upstream of APC did not rescue the growth/survival of centrinone B-treated CSC-406 organoids (data not shown).

**Fig3.**
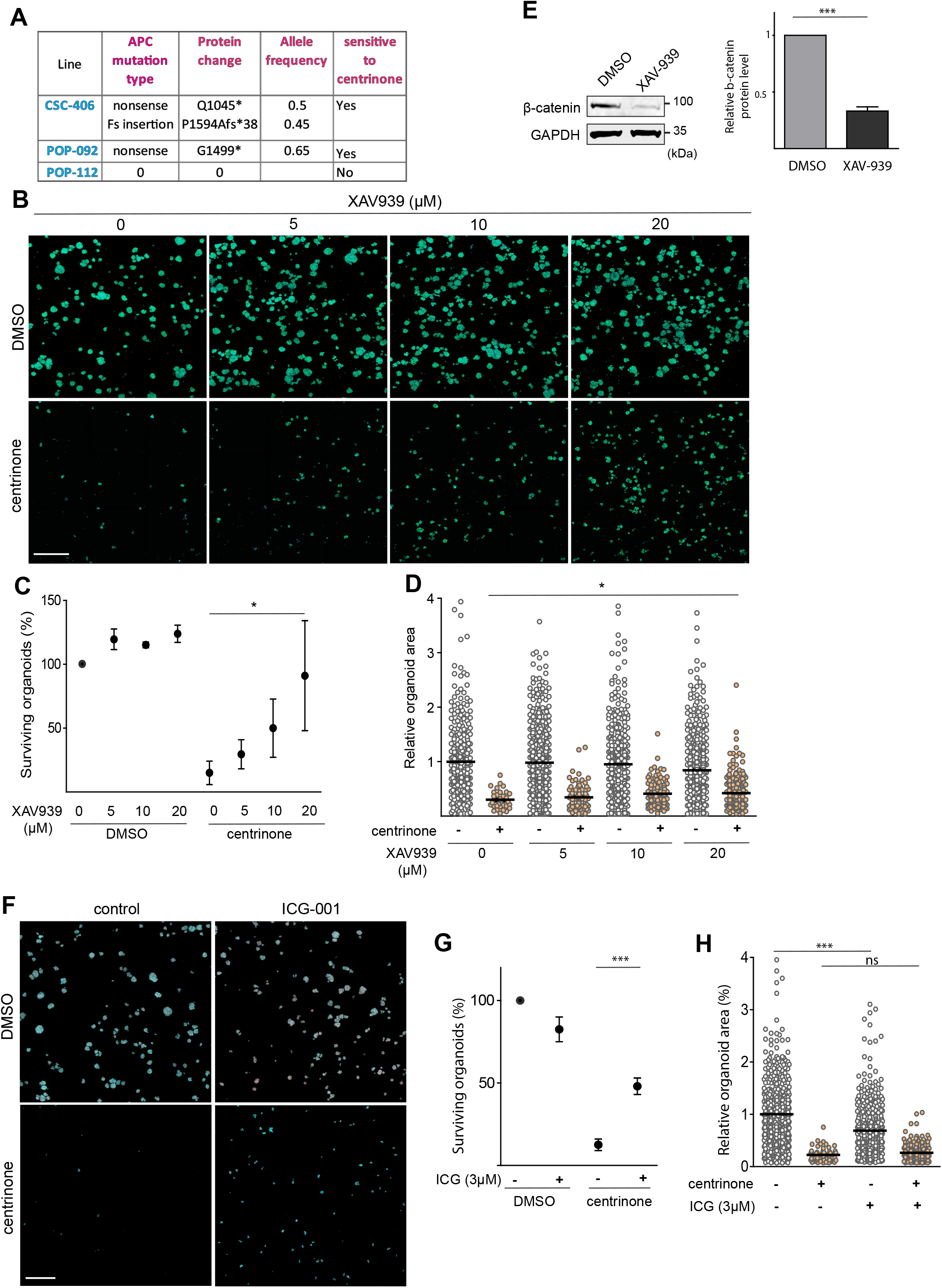
Centrosome loss-induced organoid growth defect/death is β-catenin dependent in CSC-406 cancer line. **A)** APC mutational status in the three cancer HCOs obtained via whole exome sequencing data. **B)** CSC-406 cancer organoids were grown from an equal number of single cells in the presence of DMSO or the indicated concentrations of XAV-939 and were treated with DMSO or 0.5 μM centrinone B for 8 days. Organoids were then fixed, stained with DAPI (blue) and phalloidin (green) and imaged. Merged maximum intensity projections of representative images are shown. **C)** Percentage of surviving organoids in different conditions from (B) was quantified and presented in the graph (n=3, \**P*<0.05). **D)** The areas of individual organoids (from C) were quantified in the merged maximum intensity projection images and the relative values are presented in the graph, every dot represents an organoid (n=3, \**P*<0.05). **E)** CSC-406 cancer organoids were treated with DMSO or 20 μM XAV-939 for three days, extracted from Matrigel and lysed. B-catenin and the loading control (GAPDH) protein levels were assessed using western blot analysis. Protein levels from three independent experiments were quantified and presented in the graph (n=3, \*\*\**P*<0.001). **F)** CSC-406 cancer organoids were grown from an equal number of single cells in the presence of DMSO or 3 μM ICG-001 and were treated with DMSO or 0.5 μM centrinone B for 8 days. Organoids were then fixed, stained with DAPI (blue) and phalloidin (red) and imaged. Merged maximum intensity projections of representative images are shown. **G)** Percentage of surviving organoids in different conditions from (F) was quantified and presented in the graph (n=3, \**P*<0.05). **H)** The areas of individual organoids (from F) were quantified in the merged maximum intensity projection images and the relative values are presented in the graph, every dot represents an organoid (n=3, \*\*\**P*<0.001, ns= non-significant). Scale bars are 500 μm (B, F).

To further investigate the role of the WNT pathway in the centrosome loss-induced organoid growth defect/death, we inhibited the WNT pathway in the other APC-mutant organoid line, POP-092 using the two inhibitors XAV-939 and ICG-001 as we did with the CSC-406 line. In contrast to CSC-406, XAV-939 treatment (5,10 and 20 μM) did not significantly prevent the centrinone-induced organoid growth defect/death (Fig 4A, 4B). To understand the difference between the two lines’ response to XAV-939 treatment, we determined β-catenin protein levels by western blot. We observed that the POP-092 line expressed ~90% less β-catenin protein than the CSC-406 line. Moreover, XAV-939 treatment reduced β-catenin protein levels by ~67% in CSC-406 organoids but surprisingly did not affect β-catenin protein levels in the POP-092 line (Fig 4C) probably explaining the absence of XAV-939 effect on centrinone-induced organoid growth defect/death in this line. We asked whether the POP-092 line might have lower sensitivity to the drug. Thus, we tested higher XAV-939 concentrations (40 and 80 μM) and assessed β-catenin protein levels (Fig 4D). Surprisingly, β-catenin protein levels were not affected by these concentrations. To further explore the role of β-catenin in this line, we inhibited β-catenin/TCF mediated transcription with 3 μM ICG-001 in POP-092 in the presence or absence of centrinone B (Fig 4E). Like in the CSC-406 line, ICG-001 partially prevented the centrinone-induced organoid death (~23 % rescue of the organoid survival) in POP-092 (Fig 4F) but did not rescue the organoid growth rate (Fig 4G). We noted that 3 μM ICG-001 treatment also significantly decreased the DMSO-treated POP-092 organoid growth (~25% reduction in the average organoid area) (Fig 4G). Finally, we compared β-catenin localization by immunofluorescence in the two lines. While β-catenin localized at the membrane in both lines, we found that the CSC-406 line exhibited much higher nuclear enrichment of the protein compared to POP-092 (Fig 4H).

**Fig 4.**
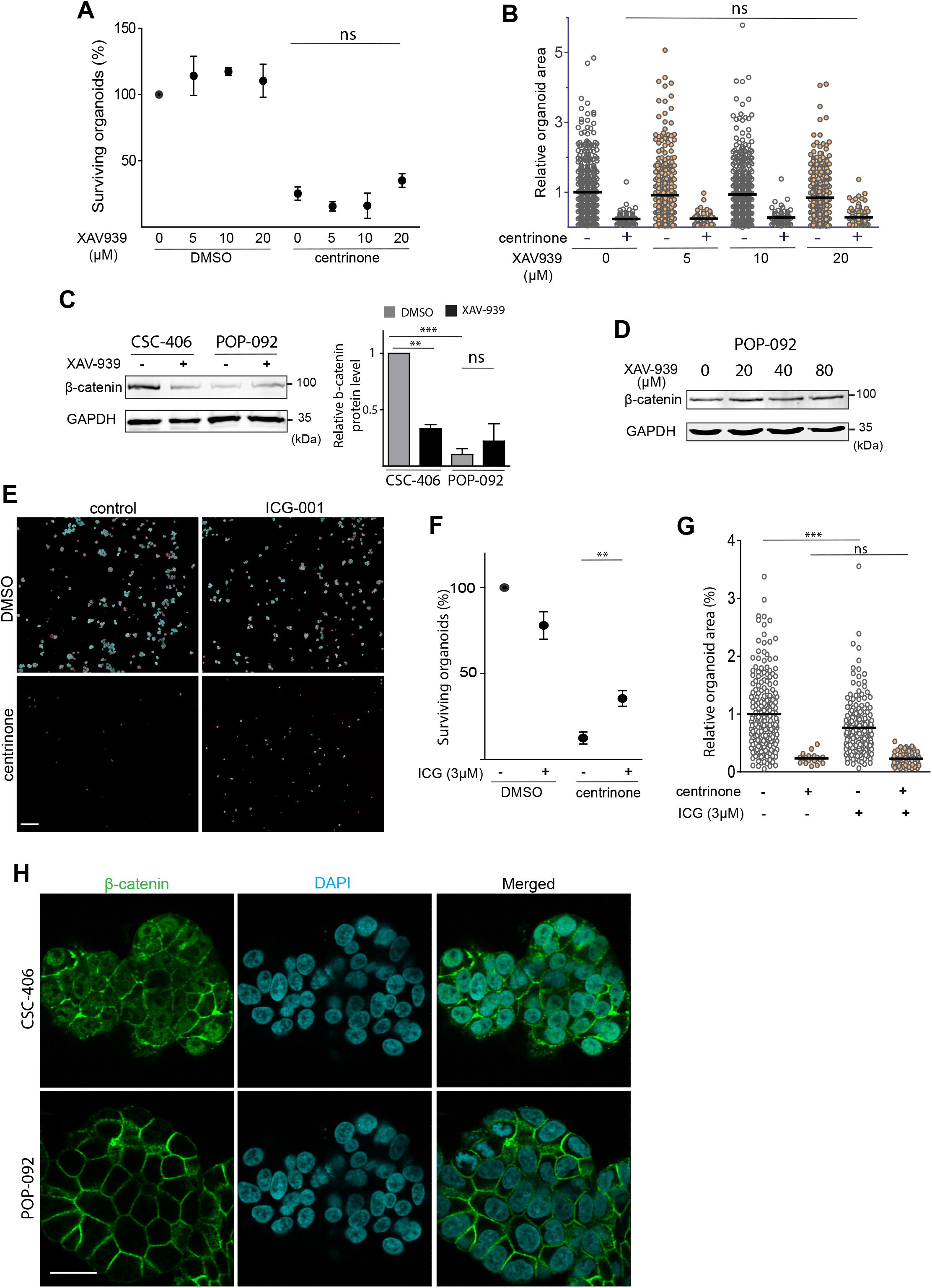
β-catenin inhibition is essential for preventing centrosome loss-induced death of POP-092 cancer organoids. **A)** POP-092 cancer HCOs were grown from an equal number of single adult stem cells in the presence of DMSO or the indicated concentrations of XAV-939 and DMSO or 0.5 μM centrinone B for 8 days. Organoids were then fixed, stained with DAP I to label nuclei and phalloidin to label actin, and imaged. Merged maximum intensity projection images were used to quantify the percentage of surviving organoids in the different conditions (n=3, ns: non-significant). **B)** The areas of individual organoids (from A) were quantified in the merged maximum intensity projection images and the relative values are presented in the graph, every dot represents an organoid (n=3, ns: non-significant). **C)** CSC-406 and POP-092 cancer HCOs were treated with DMSO or 20 μM XAV-939 for three days, extracted from Matrigel and lysed. B-catenin and the loading control (GAPDH) protein levels were assessed using western blot analysis. Protein levels from three independent experiments were quantified with Fiji and presented in the graph (n=3, \*\**P*<0.01, \*\*\**P*<0.001, ns: non-significant). **D)** POP-092 cancer HCOs were treated with DMSO or the indicated XAV-939 concentrations for three days, extracted from Matrigel and lysed. B-catenin and the loading control (GAPDH) protein levels were assessed using western blot analysis. A representative immunoblot is shown (n=2). **E)** POP-092 cancer HCOs were grown from an equal number of single cells in the presence of DMSO or 3 μM ICG-001 and were treated with DMSO or 0.5 μM centrinone B for 8 days. Organoids were then fixed and stained with DAPI to label DNA (blue) and phalloidin to label actin (Red). Representative maximum intensity projection images are shown. **F)** Percentage of surviving organoids in the different conditions tested in (E) was quantified (n=3, \**P*<0.05). **G)** organoid areas (from E) were quantified in the merged maximum intensity projection images and the relative values are presented in the graph, every dot represents an organoid (n=3, \*\*\**P*<0.001). **H)** Single focal plane images of β-catenin and DAPI staining in CSC-406 and POP-092 cancer HCOs. Note that there is more β-catenin nuclear localization in CSC-406 compared to the POP-092 line. Scale bars are 500 μm (E) and 25 μm (H).

## Discussion

Previous reports have shown that centrosome loss causes a p53-dependent growth arrest of normal human cells and that p53-mutant cancer cells continue to grow in the absence of centrosomes [1, 21]. Consistent with this, our results show that centrosome loss represses human normal colorectal organoid growth in a p53-dependent manner. It has been also shown that TRIM37 overexpression sensitizes breast tumors to centrosome loss [22]. However, our study reveals an unexpected β-catenin-mediated mechanism that controls centrosome loss-induced growth defect independently of p53 in cancer human colorectal organoids. Although p53 KO normal HCOs continued to grow without centrosomes, many abnormal nuclei phenotypes were observed including big multilobed nuclei and multi-nucleated cells most likely due to mitotic defects given the role of the centrosome in enhancing the mitotic spindle assembly efficiency [23, 24]. Despite all three cancer colorectal organoid lines carrying a non-functional mutant p53, only one line (POP-112) was resistant to centrosome loss as expected [1]. Although centrinone-resistant, this line was slightly more sensitive to low centrinone B concentrations (0.1 μM) than to the concentration needed to deplete centrosomes (0.5 μM) most likely due to centrosome amplification induced by partial PLK4 inhibition. Indeed, our previous findings demonstrate that modulating PLK4 activity in cells leads to supernumerary centrosomes at low centrinone B concentrations and centrosome loss at higher concentrations [25]. Centrinone B is >1000 fold more selective against PLK4 than

Aurora kinases AURKA and AURKB [1]. A high centrinone B concentration (10uM) results likely in a cross-inhibition of AURKA and AURKB leading to organoid growth defect of the POP-112 line independently of the centrosome.

We provide evidence that centrosome loss-induced growth defect is p53-independent in APC-mutant cancer colorectal organoids since these organoids were highly sensitive to centrinone B treatment despite harbouring a non-functional p53 [16, 17]. It has been proposed that the S240R p53 mutant (mutation in CSC-406) retains some residual (10%) transcriptional activity [26]. Given that p53 depletion did not prevent the centrinone-induced growth defect, we assume that the latter is completely p53-independent in APC-mutant colorectal cancer.

Moreover, we demonstrate that the centrosome loss-induced growth defect/death of APC-mutant cancer colorectal organoids is rather mediated by β-catenin, the downstream effector of the WNT canonical pathway. Indeed, decreasing β-catenin protein levels using XAV-939 treatment significantly prevented the centrinone-induced organoid death and partially rescued the growth of CSC-406 organoids whereas, when β-catenin protein levels were unresponsive to XAV-939 treatment in the POP-092 line, there was no significant effect of XAV-939 treatment on rescuing the centrosome loss-induced growth defect/death. XAV-939 is a Tankyrase1/2 inhibitor that stabilizes Axin, a central scaffolding protein in the β-catenin destruction complex leading to β-catenin phosphorylation, ubiquitination, and degradation [19, 27]. It is not clear why XAV-939 treatment in the POP-092 line does not affect β-catenin protein levels. Axin protein levels are regulated through Tankyrase-mediated PARsylation and ubiquitination targeting it to proteasome degradation [19]. The unresponsiveness of the POP-092 line to XAV-939 may be due to an altered signaling mechanism upstream of Axin (PARsylation and ubiquitination) or a failure to enhance the β-catenin destruction complex assembly downstream of Axin. Exploring Axin protein levels in response to XAV-939 treatment will help address this question.

The role of β-catenin in mediating centrosome loss-induced cancer organoid death was confirmed in both APC-mutant lines (CSC-406 and POP-092) since the inhibition of β-catenin/TCF mediated transcription using ICG-001 partially prevented centrinone-induced death of both lines, although the effect on organoid growth rate was insignificant compared to XAV-939, most probably due to ICG-001 cell toxicity effect [20]. Finding the balance between an effective and nontoxic ICG-001 concentration was challenging. This might explain the better organoid growth rescue obtained with XAV-939 which is better tolerated due to the non-complete inhibition of β-catenin. Comparing the APC-mutant cancer organoids shows that APC is more mutated in CSC-406 (nonsense Q1045*, allele frequency 0.5 / Fs insertion P1594Afs*38, allele frequency 0.45) compared to POP-092 (nonsense G1499*, allele frequency 0.65) indicating a higher WNT pathway activity which correlates with higher β-catenin protein levels and its increased nuclear enrichment in CSC-406 compared to POP-092 line. This may also explain why the CSC-406 line is slightly more sensitive to centrosome loss and responds better to β-catenin inhibition since it is more dependent on WNT/ β-catenin pathway compared to the POP-092 line.

In summary, our study identifies WNT/ β-catenin as a new signaling pathway that mediates centrosome loss-induced growth defect and death in APC-mutant cancer colorectal organoids independently of the canonical p53 pathway. Therefore, targeting the centrosome may be a good strategy to combat colorectal cancer with non-functional p53 and hyperactive WNT/ β-catenin pathway.

## Materials and methods

### Reagents

Centrinone B *(Tocris Bioscience* #5690/10), Alexa Fluor^™^ 488 Phalloidin (Invitrogen #A12379), Alexa Fluor^™^ 546 Phalloidin (Invitrogen #A22283), DAPI (Invitrogen #21490), XAV-939 (Cayman Chemical #13596-10), ICG-001 (Cayman Chemical #16257-5), Nutlin-3a *(Cayman Chemical* #10004372).

### Antibodies

p53 (Santa Cruz #SC126, WB: 0.2 μg/ml, IF: 0.8 μg/ml), CEP192 (Bethyl #A302-324A, IF: 4 μg/ml), pericentrin (Abcam #ab4448, IF: 1 μg/ml), GAPDH (Sigma #G9545, WB: 1 μg/ml), β-actin (Sigma #A5316, WB: 1 μg/ml), β-catenin (Santa Cruz # sc-7963, WB: 0.2 μg/ml, IF: 1 μg/ml), anti-mouse Alexa Fluor 488 (ThermoFisher # A-21202, IF: 0.5 μg/ml), anti-rabbit Alexa Fluor 488 (ThermoFisher # A-21206, IF: 0.5 μg/ml).

### Human organoid lines

Normal and cancer Human Colorectal Organoids (HCOs) were obtained with informed patient consent from UHN Princess Margaret Living Biobank (PMLB, Toronto, Canada). All studies were approved by the Mount Sinai Research Ethics Board (MSH REB Study # 18-0101-E). Three cancer organoid lines derived from three different patients: CSC-406 (female, 69 years old, colorectal adenocarcinoma), POP-092 (female, 46 years old, colorectal adenocarcinoma) and POP-112 (male, 74 years old, colorectal adenocarcinoma) were used in our study. The normal organoid line and CSC-406 cancer line were derived from the same patient. To make the p53 KO normal and cancer lines, we produced lentiviruses in HEK293T cells using TLCV2 lentiviral vector (Addgene #87360) expressing a Tet-inducible CRISPR-Cas9 protein, in which we cloned a gRNA targeting the genomic p53 sequence: TATCTGAGCAGCGCTCATGG as described by the cloning protocol provided by Addgene (#87360). Organoids were broken up into single cells using a 40 min incubation (37°c) with TrypLE ^™^ express (Gibco^™^ #12605-028) followed by filtering through a 30 μM cell strainer (PluriSelect #43-50030-03). Single cells were then infected with the lentivirus in the complete growth media on a Matrigel layer. The day after, adult stem cells were attached to the Matrigel. Media was removed, another layer of Matrigel was added and fresh growth media containing puromycin (3 μg/ml) was supplemented. After 2 days, CAS9 protein expression was induced using 1μg/ml Tetracycline. After 3 days, p53 KO organoids were enriched by Nutlin-3a treatment (10 μM for 2 weeks) [14]. p53 Knockout efficiency was verified using TIDE analysis, immunofluorescence, and Western Blot.

### TIDE analysis

Genomic DNA was extracted from control and p53 KO organoid lines (PureLink^™^ Genomic DNA Mini Kit, Invitrogen^™^#K182001) and a PCR was performed to amplify the genomic region targeted by the p53 guide using the primers: CTCAACAAGATGTTTTGCCAAC (Forward) and ACTCGGATAAGATGCTGAGGAG (Reverse). The PCR products were then Sanger sequenced using the Forward primer. Analysis of the two resulting raw sequencing files using the TIDE web tool (https://tide.nki.nl/) identifies the indels and their frequencies, giving a knockout efficiency score [15].

### Organoid culture

Normal and cancer Human Colorectal Organoids (HCOs) were obtained from UHN Princess Margaret Living Biobank (Toronto, Canada). HCOs were embedded in Matrigel domes and maintained in growth media composed of: Advanced DMEM/F-12 (Gibco^™^ #12634010), 100 U/ml penicillin/streptomycin (Gibco^™^ #15140-122), 10 mM HEPES (Gibco^™^ #15630-056), 2 mM GlutaMAX^™^ (Gibco^™^ #35050), 1.25 mM *N*-acetyl-cysteine (Sigma-Aldrich # A9165-5G), 1× B27 supplement (Gibco^™^ #17504-044), 10 nM gastrin (Sigma #G9145), 50 ng/ml mouse EGF (Gibco^™^ #PMG8041), 100 ng/ml mouse Noggin (Peprotech #250-38), 0.5 μM TGFb type I Receptor inhibitor A83-01 (Tocris #2939). In addition, Normal HCO media was supplemented with 40% WNT-3a conditioned media, 10% R-spondin conditioned media (UHN Princess Margaret Living Biobank, Toronto, Canada), 2.5 μM CHIR 99021 (Tocris #4423) and 10 μM p38 MAPK inhibitor, SB202190 (Sigma-Aldrich # S7067) while the cancer HCO media contained 10 μM of the rock inhibitor Y27632 (MedChem Express # HY-10583). Media was changed every 2 to 3 days and organoids were passaged every 7-9 days. Normal HCOs were broken up using enzymatic digestion using TrypLE ^™^ express for 1-2 min followed by mechanical disruption (rigorous 10 times up and down pipetting) while only the enzymatic method (10 min TrypLE ^™^ Express) was used for passaging the cancer HCOs. In experiments when starting organoid cultures (normal and cancer) from single cells is needed, a 30-45 min TrypLE ^™^ Express incubation at 37°c was performed, followed by cell filtering through a 30 μM cell strainer (pluriSelect #43-50030-03).

### Immunofluorescence

Organoids were embedded in Matrigel and grown on Nunc^™^ Lab-Tek^™^ II Chambered Coverglasses (Thermoscientific^™^ # 155360) for the indicated days. After cold PBS wash on ice (10 min), organoids were fixed with cold 4% paraformaldehyde (at 4°c for 20-25 min). Cold PFA causes Matrigel meltdown allowing organoids to be fixed to the bottom of Chambered Coverglasses. After 1h permeabilization with 0.5% cold triton and 2-3h blocking with 1.5% BSA solution containing 0.1% triton, organoids were incubated with the primary antibodies in the blocking solution overnight at 4°c. The day after, organoids were washed three times with PBS and incubated with secondary antibodies coupled with Alexa Fluor-488 at 0.5 μg/ml concentration for 2-3h. DNA was stained with DAPI and F-actin with Phalloidin (1/500). Organoids were PBS washed, then imaged in the imaging solution (0.7 mM N-acetyl cysteine, PH~7.4).

### Imaging and image analysis

Fixed and stained organoids were imaged in Nunc^™^ Lab-Tek^™^ II Chambered Coverglasses (Thermoscientific^™^ # 155360) using Nikon A1 HD25/A1R HD25 confocal microscope equipped with NIS-Elements controller. 20X and 40X water immersion objectives were used to take z-stack images. To quantify organoid size and number, merged maximum intensity projections (DAPI and Phalloidin) of large images were processed in Fiji software as follows: image > type > 16-bit > adjust threshold > analyze particles. Fiji presents then the count and the size of all areas in the image. Each area represents an organoid.

### Western blotting

Organoids were extracted from Matrigel using TrypLE ^™^ Express (20 min) and then lysed in lysis buffer (50 mM Tris-HCl, 1% NP-40, 10% glycerol, 140 mM NaCl, 5 mM MgCl2, 20 mM NaF, 1 mM NaPPi and 1 mM orthovanadate, pH 7.4) complemented with protease inhibitor cocktail (Roche #11697498001). Total soluble proteins were measured using BCA Protein Assay Kit (Thermo Scientific #23227), then denatured in SDS sample buffer. Equal protein amounts were run on polyacrylamide gels and transferred onto PVDF membranes. After milk blocking, membranes were incubated with the specific primary antibodies in the blocking solution overnight. After TBS-Tween wash, membranes were incubated with the fluorescently labeled secondary antibodies and proteins were detected using Licor Odyssey CLx (LI-COR Biosciences - U.S). The digital images obtained were quantified using Fiji software.

### Statistical analysis

Statistical analysis was performed using a two-tailed t-test, one-way or two-way ANOVA analysis of variance followed by a Bonferroni’s multiple comparison test using GraphPad Prism (version 8, https://www.graphpad.com/scientific-software/prism/).

## Acknowledgements

We thank Mardi Fink from Dr. Jeff Wrana’s lab at the Lunenfeld-Tanenbaum Research Institute for her help with establishing colorectal organoid culture in our lab.

## Author contributions

MB and LP designed the project, MB performed all the experiments, analyzed the data, and wrote the manuscript with editing from LP. NR and MST established the UHN organoid core, generated the normal and colorectal cancer lines used in this study and Whole Exome Sequencing (WES) data. This work was funded by grants from the Krembil Foundation and the CCSRI to LP.

